# A Mean Field Theory for Pulse-Coupled Oscillators based on the Spike Time Response Curve

**DOI:** 10.1101/2025.01.09.632091

**Authors:** Carmen C Canavier

## Abstract

A mean field method for pulse-coupled oscillators with delays used a self-connected oscillator to represent a synchronous cluster of N-1 oscillators and a single oscillator assumed to be perturbed from the cluster. A periodic train of biexponential conductance input was divided into a tonic and a phasic component representing the mean field input. A single cycle of the phasic conductance from the cluster was applied to the single oscillator embedded in the tonic component at different phases to measure the change in the cycle length which the perturbation was initiated, that is, the first order phase response curve (PRC), and the second order PRC in the following cycle. A homogeneous network of 100 biophysically calibrated inhibitory interneurons with either shunting or hyperpolarizing inhibition tested the predictive power of the method. A self-consistency criterion predicted the oscillation frequency of the network from the PRCs as a function of the synaptic delay. The major determinant of the stability of synchrony was the sign of the slope of the first order PRC of the single oscillator in response to an input from the self-connected cluster at a phase corresponding to the delay value. For most short delays, first order PRCs correctly predicted the frequency and stability of simulated network activity. However, considering the second order PRC improved the frequency prediction and resolved an incorrect prediction of stability of global synchrony at delays close to the free running period of single neurons in which a discontinuity in the PRC precluded existence of 1:1 self-locking.

## Introduction

Synchronization tendencies in coupled oscillator networks are often studied by applying control theory under the assumption of weak coupling [1,2] that each oscillator remains very near its unperturbed trajectory, called a limit cycle, or under the assumption of pulsatile coupling [3–5] that each oscillator returns to its unperturbed trajectory before the next input is received. Connectivity between neurons is generally mediated by a synaptic conductance waveform triggered by the arrival of a presynaptic spike. There is a substantial literature on pulse-coupled oscillators which was reviewed in [4], with many advances in pulse-coupled oscillators with delayed coupling [6–12] occurring since that review. Here we address synchronous fast oscillations in which the period of the network oscillation is not necessarily long compared to the duration of the conductance waveform that mediates the coupling. In that case, there can be a substantial summation of effects from previous inputs, leading to a tonic component of the coupling. The tonic component distorts the original limit cycle, breaking both weak and pulsatile coupling assumptions and requiring a different formalism which is presented here.

Examples of fast neural oscillations include ripples [13] and fast gamma oscillations [14]. Theta-nested gamma oscillations are thought to play a fundamental role in the initial encoding and retrieval of memories, and sharp wave ripples are thought to play a fundamental role in consolidation of memory representations [15]. Theta-nested fast gamma (65-140 Hz) in the medial entorhinal cortex is hypothesized to convey information regarding current sensory representations [14]. Ripple frequencies are variously defined, but their lower bound is 100 Hz [16] to 140 Hz [13], and they extend up to 200 Hz and beyond. The role of inhibition in ripple synchronization has been debated [17,18]. However, numerous studies show that parvalbumin positive fast spiking interneurons (PV-INs) are phase-locked to ripple cycles and often fire at ripple frequency both *in vivo* [19–26] and *in vitro* [27–31]. Moreover, blocking GABA_A_ receptors abolishes ripples [32]. PV-INs have also been shown to mediate gamma synchrony [33–35]. In some cases, gamma oscillations can be supported by inhibitory interneurons alone [36–38], although there is frequently an interplay between inhibitory and excitatory cells [39]. PV-INs are thought to be specialized for rapid signaling [40]. Indeed, our group has shown that the synapses between PV-INs are fast, with delays on the order of ∼0.8 ms, a rising exponential time constant of ∼0.3 ms and a falling exponential time constant of ∼2 ms [41]. Despite these rapid kinetics, in a network oscillation at 200 Hz, the biexponential waveform lasts several cycles, providing biological inspiration for the novel framework presented herein.

We assume a synchronous population in which each neuron receives identical tonic and phasic components that result from a train of inputs. The input from the network is received with a delay that incorporates conduction and synaptic delays. The basic idea is to represent a synchronous neural network with a self-connected oscillator, perturb a single neuron, and apply phase resetting theory to determine which delays produce stable global synchrony and to predict the frequency of that synchronous oscillation. Ultimately, the insights gained into the conditions that support synchronization may enable therapeutic strategies to enhance synchrony that underlies cognitive function or to disrupt pathological synchrony.

For illustrative purposes, we chose a recent model [42] of a network of 100 parvalbumin positive inhibitory interneurons with properties carefully calibrated to match data from layer 2/3 in the medial entorhinal cortex [41]. For ease of analysis, we used the homogeneous version of the network, in which each neuron is identical, receives identical simulated excitation, and receives exactly N=36 synaptic inputs from the other interneurons in the network. Thus the perturbed neuron receives N = 36 inputs from the remaining self-connected cluster, and the self-connected cluster receives N-1=35 delayed inputs from itself plus one input from the perturbed oscillator. These numbers are specific to our example, and should be adjusted to fit the particular application.

## Results: Theoretical Framework

### 2.1 The mean field produced by a train of biexponential conductance waveform

The biexponential synapse [43] is often used to approximate the conductance waveform that results when the arrival of an action potential (spike) at a presynaptic terminal of an ionotropic synapse initiates a release of a neurotransmitter. The neurotransmitter diffuses across the synaptic cleft and binds to postsynaptic receptors that directly open ligand-gated ion channels. As the neurotransmitter diffuses away from the cleft, channels unbind from the ligand and close. These processes cause an initial increase in synaptic conductance that then decays. Multiplying a rising exponential by a decaying exponential simulates the resultant conductance waveform: 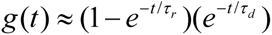, where the subscript *r* indicates a rising and the subscript *d* indicates a decaying exponential.

The biexponential synapse can be also described by a difference of exponential terms by multiplying the rising and falling exponentials to obtain 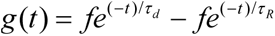, where, *t* is the time elapsed since the arrival of a presynaptic spike at *t*=0, the factor to normalize the peak of a single biexponential synaptic conductance waveform to 1 is 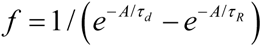 and A is the time at which the maximum value of the biexponential occurs: 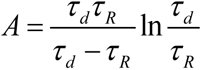. Note that in this form, the decay time constant τ_d_ is preserved, but only when τ_d_ >>τ_R_ does the rise time constant τ_R_ approximate the true rise time τ_r._ If a train of such biexponential synapses are activated with period *P_F_* starting with a spike arriving at *t*=*t_0_* and *t_0_+nT* is the arrival time of the most recent presynaptic spike, then the following expression applies:

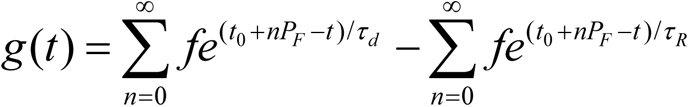

Rearranging

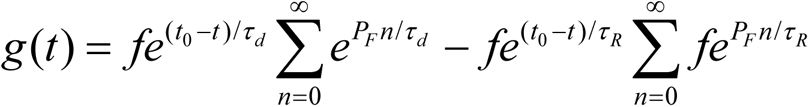

We can use the formula for the sum of an infinite power series 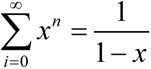 to get the time course of the conductance after an infinite number of activations:

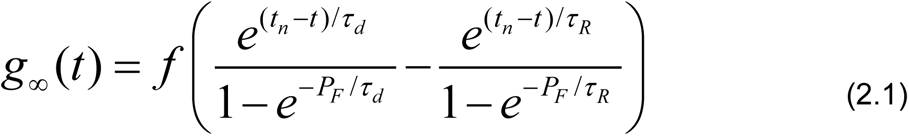

for sufficiently large values of *n* because the waveform after each spike received at time *t_n_* is identical. The minimum value of the response to an infinite train occurs at the arrival times *t_n_*, again for sufficiently large:

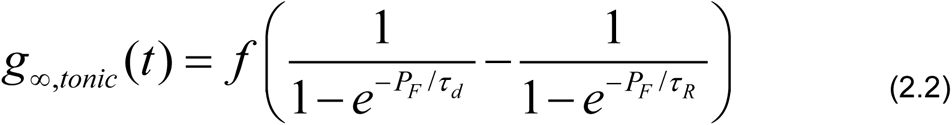

This is the tonic component we propose to apply to the uncoupled oscillator to form a new limit cycle oscillator. The tonic component accounts for the effects of all previous inputs from a synchronous train of biexponential inputs that comprise the mean field received by each individual oscillator in a synchronous network. To determine the response of each individual oscillator to the phasic portion of the mean field input, we subtract the tonic component in Eq. 2.2 from the steady-state conductance waveform given Eq. 2.1 to obtain the phasic component

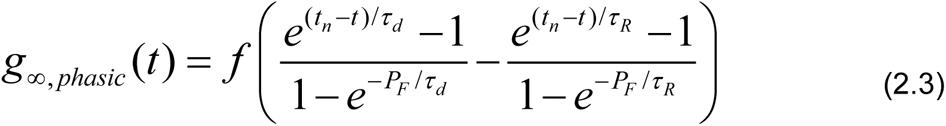

where *t_n_* is the time at which the *n*th perturbation is applied. Each spike in a periodic train evokes a single biexponential conductance waveform with a peak at 1. The peak of the train (Figure 1) can be larger than 1 because of the accumulated contributions of previous events in the train. The minimum of the waveform is the tonic component (horizontal dashed line) and the phasic component is located between the two vertical dashed lines, here for a train applied every 5 ms and extending infinitely back into the past.

**Figure 1.**
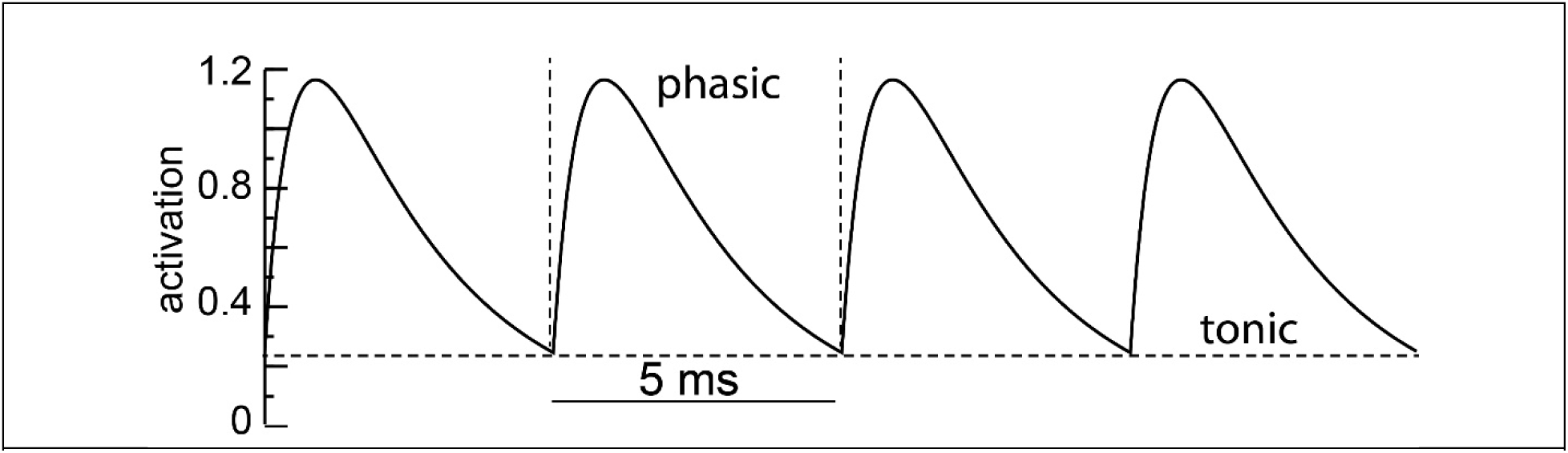
Biexponential Conductance Train. A train of biexponential conductances at 200 Hz with τ_R_ = 0.3 ms and τ_d_=2.0 ms.

### 2.2 Mean Field Phase Response Curve

To determine the response of an individual oscillator to a phase shift of the event times *t_n_,* at which an input is received from the network relative to the spikes emitted by the individual oscillator, we must first transform the uncoupled individual oscillator with intrinsic period *P_i_* to an oscillator embedded in the tonic portion of the mean field with a new baseline period *P_E_* due to the embedding. This requires adding the tonic conductance from Eq. 2.2, at an assumed forcing period P_F_, to a synaptic current, which could be an inhibitory GABAergic synaptic current or an excitatory AMPA synaptic current, with the appropriate reversal potential to provide the driving force for the synaptic current. Experimentally, one could use the dynamic clamp [44,45] to inject a virtual conductance. The resultant embedded free running period is P_E_ in Figure 2A, which illustrates the computation of the mean field PRC for two values of the inhibitory reversal potential, indicated by the horizontal dashed lines superimposed on the voltage traces. A value of -55 mV is shunting because it is within the range of values traversed during the interspike interval, but remains below the action potential threshold whereas a value of -75 MV is hyperpolarizing because it is more hyperpolarized that the values traversed during the interspike interval. The same number of cycles are shown for each case; however, a close examination of the scale bars reveals that the embedded period is shorter for shunting compared to hyperpolarizing inhibition.

**Figure 2.**
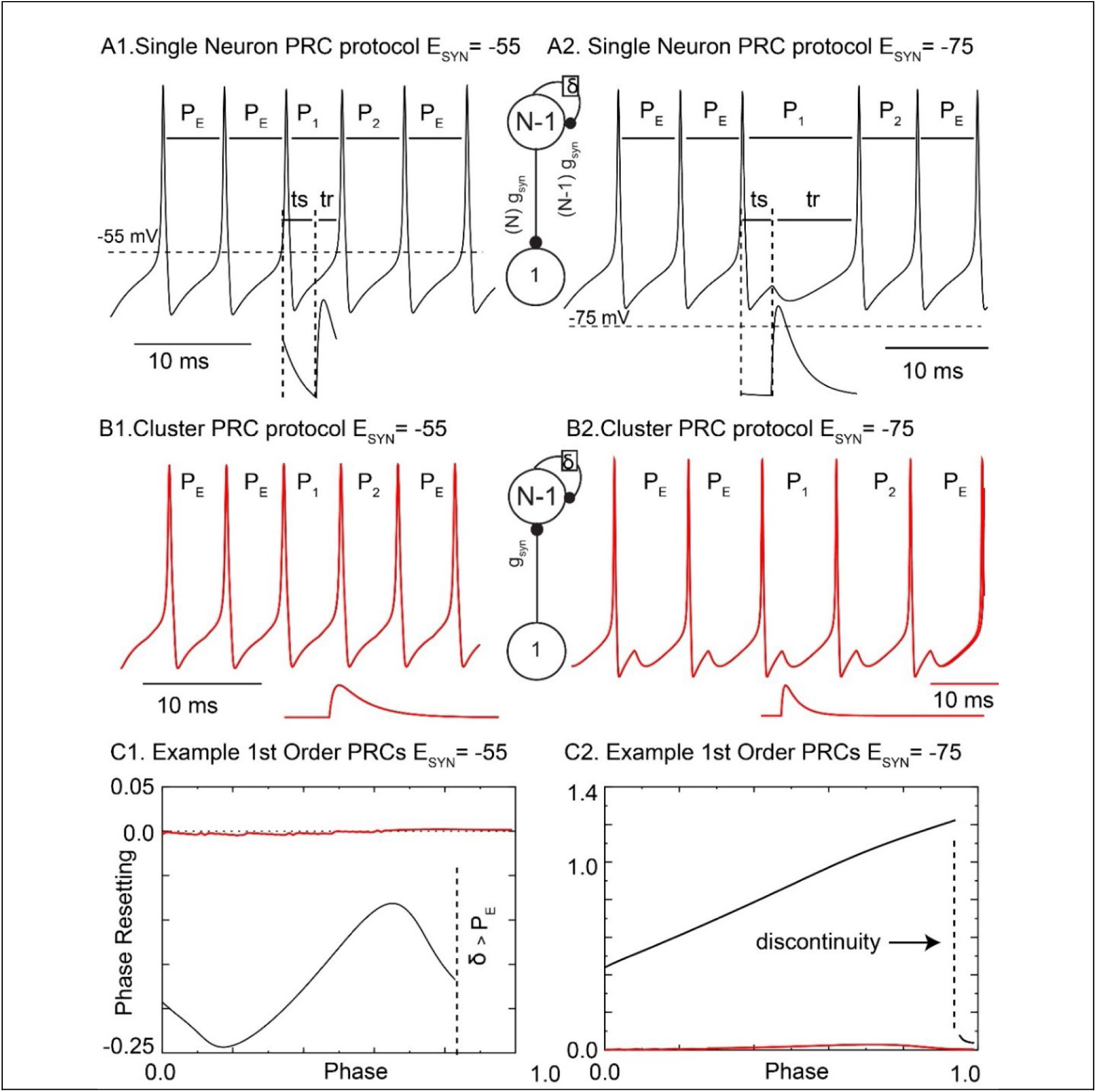
Phase resetting curves for perturbed single neuron and self-connected cluster. **Left:** Shunting Inhibition. **Right:** Hyperpolarizing inhibition. **A.** Mean field Analysis: Voltage traces (top) of a single neuron and phasic conductance waveforms (bottom) received from the cluster (see inset) for a 3 ms delay to the peak of the input. Interval labels are explained in the text. We assume P_E_ is restored after P_2_. **B.** Spike Time Response Curve: Voltage traces (top) of a self-connected neuron and biexponential conductance waveforms (bottom, not to scale because the input from the single neuron is much smaller than that from the cluster) from the single neuron (see inset) for a 3 ms delay to the peak of the input. **C**. Representative Phase Response Curves (PRCs) for first order resetting of the single neuron (black) and the self-connected cluster (red) as computed in A and B respectively. In **C1** the vertical dashed line indicates that the PRC cannot be measured as explained in the text whereas in **C2** it indicates a discontinuity.

The phasic effect of the network is determined by applying one cycle of the phasic conductance waveform in the cycle labeled P_1_. The trough representing the timing of the input is shifted relative to a spike emitted by the individual embedded oscillator to determine the effect of delays on the resultant phase-locked network period *P_N_*. This is equivalent to a sudden change in phase of the perturbed single oscillator (inset in Figure 2A), relative to the synchronous activity of the ensemble of N-1 neurons, due to noise. The interval between a spike emitted by the individual embedded oscillator and the trough corresponding to the receipt of an input is called the stimulus interval *ts*, which for a synchronous network is equal to the delay δ. We define phase φ as ranging from 0 at the time the membrane potential crosses the threshold for spike detection (here set at -30 mV) to 1 at the time the next free running spike is emitted. Thus, for a given delay δ, an input in a 1:1 globally synchronous mode is received at a phase φ= δ/P_E_. For a mean field PRC, prior to the trough, the phasic part of the synaptic conductance waveform results from an input prior to the current cycle that occurred at time =0, P_F_-ts ms prior to the spike at phase 0 and is given by

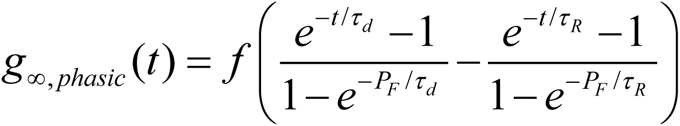

at times t =[P_F_-ts , P_F_). After the trough, the phasic synaptic conductance waveform is due to the input applied at the trough and is given by the following equation at times t = [P_F_,2P_F_-ts].

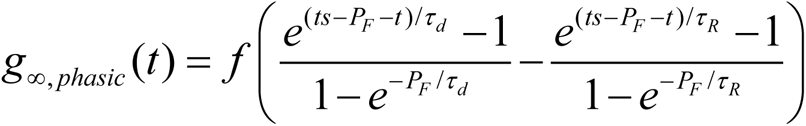

The examples in Fig. 2A were chosen such that the assumed forcing period produces a phasic synaptic conduction waveform (bottom traces) the same length as P_1_, the perturbed cycle, as explained in the text accompanying the next figure. The effects on both the cycle (P_1_) containing the perturbation (first order resetting) and the subsequent cycle (P_2_, second order resetting [46,47]) can be included in mean field theory. First order resetting is defined as the normalized change in P_1_ cycle length due to the perturbation: f_1_(φ)= (P_1_-P_E_)/P_E_. Similarly, second order resetting is defined as f_2_(φ)= (P_2_-P_E_)/P_E_. However, we will initially consider only first order resetting, since in many cases it is sufficient and second order resetting can be ignored. The interval between the trough of the phasic component and the emission of the next spike is called the response interval *t_r_*. This interval is determined by the interval remaining in the unperturbed cycle until the next spike plus the first order phase resetting: *t_r_= P_E_(1-φ +f_1_(φ)).* Figure 2C (black traces) gives example mean field first order phase resetting curves. If a second spike is emitted by the embedded oscillator prior to the trough, the phase response cannot be calculated, indicated by the “δ> P_E_” region to the right of the vertical dashed line in Fig. 2C1. The vertical dashed line in Fig. 2C2 indicates a discontinuity commonly observed in phase responses to strong hyperpolarizing inhibition[48–52]. This discontinuity arises at late phases because the bulk of the area under the conductance waveform occurs in P_2_ rather than P_1_, and the inhibition in P1 is not sufficient to prevent the next spike, thus the abrupt decrease in the magnitude of the phase delay.

Figure 2B shows the responses of the self-connected cluster to a inputs from a single perturbed neuron for the same two values of E_SYN_ as in Fig. 2A. In this case, we neglected the effect of the single neuron on the embedded period and simply used the free-running period of the self-connected cluster as P_E_. We used the unaltered biexponential waveform as the perturbation to the cluster, called a spike time response method [53] rather than a mean field version. This is a potential source of error, but the results obtained with this simplification justified the assumptions in this case, as explained below. The effects of the self-inputs to the cluster are obvious in Fig. 2B2 but not in Fig. 2B1. The resultant first order PRCs of the self-connected cluster (red traces in Fig. 2C) are negligible compared to those of the single neuron (black traces).

The predictive power of the PRC lies in the assumption that the stimulus and response intervals measured in the “open-loop” configuration shown in the insets of Fig. 2A and B are preserved in the “closed-loop” configuration shown in the inset to the left of Fig. 3A. The assumed perturbed pattern from synchrony is illustrated in Fig. 3A1 and steady state synchrony with short delays is given in Fig. 3A2. The critical feature of this firing pattern is that it takes k=2 network periods [6] for a spike one oscillator to affect the timing of a spike in the same oscillator via the feedback loop through the other oscillator. The first red arrow points from the first spike in the single oscillator to the spike in the other oscillator that the delayed input affects, and the second arrow points from the perturbed spike to the spike in the single oscillator whose timing is affected by the delayed input from that spike. This completes the feedback loop. The firing pattern in Fig. 3A1 produces a discrete map of the firing intervals that allows the derivation of the stability criterion given in the next section. Givin the perturbation in spike times Δ[n], the stimulus intervals are δ + Δ[n] and δ – Δ[n] respectively. The response intervals can be predicted using the PRC as described above *tr= P_E_(1-φ +f_1_(φ))*. The new value of the perturbation in spike timing can be calculated as Δ[n+1] = Δ[n] + ts_1_[n]+tr_1_[n]-ts_N-1_[n]+tr_N-1_[n]. These steps can be repeated indefinitely; for a stable mode Δ[n] goes to 0 as n goes to ∞. If global synchrony exists, the intervals are described by Fig. 3A2 in which the stimulus interval is exactly equal to the delay. For a one-to-one locking between the forcing from the network and spikes in the embedded oscillator to exist at a specific delay δ, one existence criterion is *P_F_ =P_E_(1+f_1_(δ/P_E_))*, meaning that the forcing period produces an identical network period as predicted by phase resetting theory (for now neglecting second order resetting). This criterion lends itself to a graphical solution in which the network period is plotted for each value P_F_ and any intersection of the resultant curve with the diagonal line y=x line determines the network period of a synchronous solution with P_N_ = P_F_ at a given delay as illustrated in Figure 3B1 and B2 for the two cases illustrated in Fig. 2A and B, with predicted network periods of about 5 and 11 ms respectively at delays equal to 3 ms.

**Figure 3.**
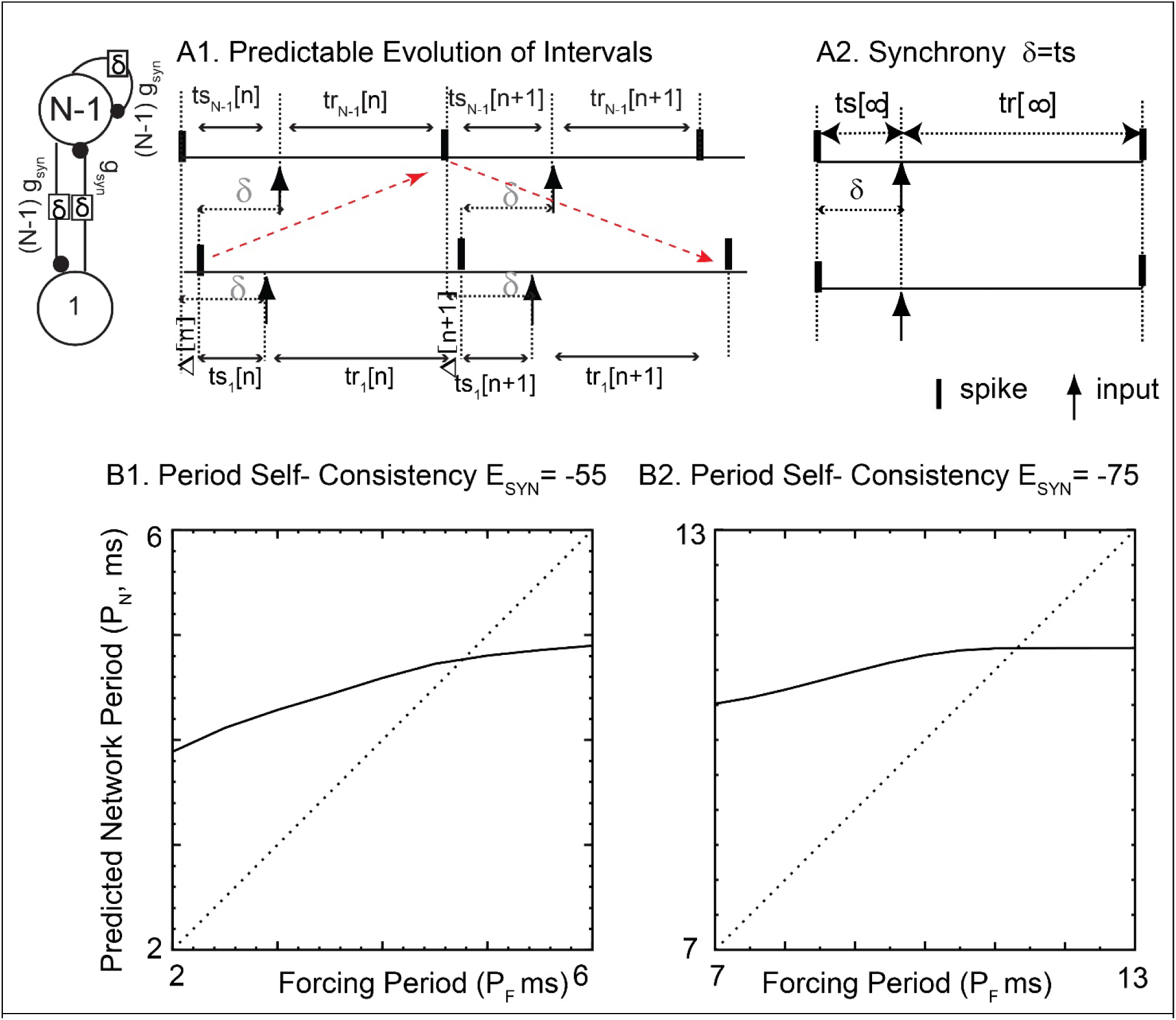
Existence of Global Synchrony. A. Closed loop interval maps. A1. Map of the intervals in an alternating mode with delays for a single neuron perturbed from the cluster by interval Δ. A2. Steady state intervals in global synchrony. B. Existence criterion for Network Period. B1. Shunting inhibition. B2. Hyperpolarizing inhibition.

### 2.3 Stability of synchrony

Recently, results for two pulse-coupled oscillators with delays [6] were extended to global synchrony in a homogeneous network. Here we consider firing patterns in which the firing of one oscillator affects a subsequent firing in the same oscillator via the feedback pathway through the other oscillator after exactly two network periods (Fig. 3A2), and specifically a perturbation from global synchrony. The following criterion is necessary for synchrony to be a stable solution” for k=2 in the terminology in the study that originally derived it [7]:

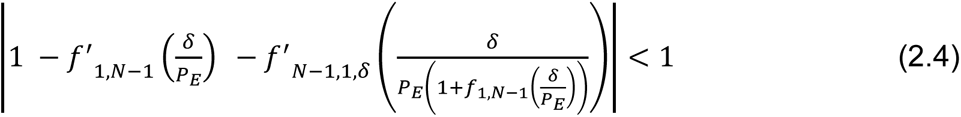

We replaced the intrinsic period in the original criterion with the embedded period of the single neuron to generalize to the mean field case. The criterion depends upon the slopes of two first order phase resetting curves at their respective locking phases at a given delay: the response of the single neuron to a population spike in a cluster containing N-1 neurons, indicated by the subscript 1, N-1 and an additional phase resetting curve we have not previously addressed, indicated by three subscripts N-1,1, δ. The latter subscripts indicate the response of the remaining N-1 embedded neurons, which are self-connected with a delay δ, to a spike in the individual embedded neuron. In order to get the phase of the self-connected cluster, the delay is normalized by the period of the self-connected cluster, which can be computed from the PRC of the single neuron. For our example case, the slopes of the first order PRC for the self-connected cluster were so small (red traces in Fig. 2C) that they were neglected, and the slope of the first order PRC for the single neuron dominated the stability criterion. If that slope was positive (but less than two) the absolute value of the eigenvalue in Eq.2.4 was less than one predicting stable 1:1 global synchrony, whereas if it was negative the absolute value was greater than one predicting that synchrony was unstable.

## 3 Results: Testing the theory in a simulated network of homogeneous neurons

We ran simulations of the 100 neuron homogeneous network using publicly available code downloaded from https://modeldb.science/267338. For all 1:1 synchronous modes detected, we plotted the frequency as the open black circles in the top panels of Fig. 4A and B for shunting and hyperpolarizing inhibition respectively. The filled red circles show the predicted frequency for stable 1:1 synchronous modes predicted according the existence and stability criteria described above at each value of the network delay up to the intrinsic period of the unperturbed single neuron, about 6 ms. The bottom panels show the slope of the first order mean field PRC for the single neuron. The PRC must be measured at each feedback delay value, then the slope is taken at the phase (delay normalized by embedded period) corresponding to the same delay. Two regions of the PRC for shunting inhibition in the bottom panel of Figure 4 A have a negative slope, hence synchrony is predicted to be unstable in those regions. Accordingly, no simulations of global 1:1 synchrony were found in the simulations, with a couple of exceptions near the predicted boundary at a delay of 3.2 ms. In a narrow range near 3.1 ms, multistability between synchrony and an asynchronous mode was identified by using different initializations of the system. There was a slight systematic overestimation of the observed frequency (filled red versus open black circles). In Fig. 4B, there was a slight systematic underestimation of the observed frequency. There was general agreement that positive values of the slope predicted synchrony. However, there was an anomaly at late phases (delays close to the values of the free running intrinsic period of the single neuron). Synchrony at values of the delay between 4.6 and 4.9 ms was increasingly difficult to detect due to multistability with an unusual mode described in the next figure. Whereas the stability analysis predicted synchrony should not become unstable until the delays reached a value of 5.7 ms, synchrony was not observed in the simulations beyond a delay value of 4.9 ms. This forced us to re-evaluate our stability criterion to include second order resetting of the single neuron.

**Figure 4.**
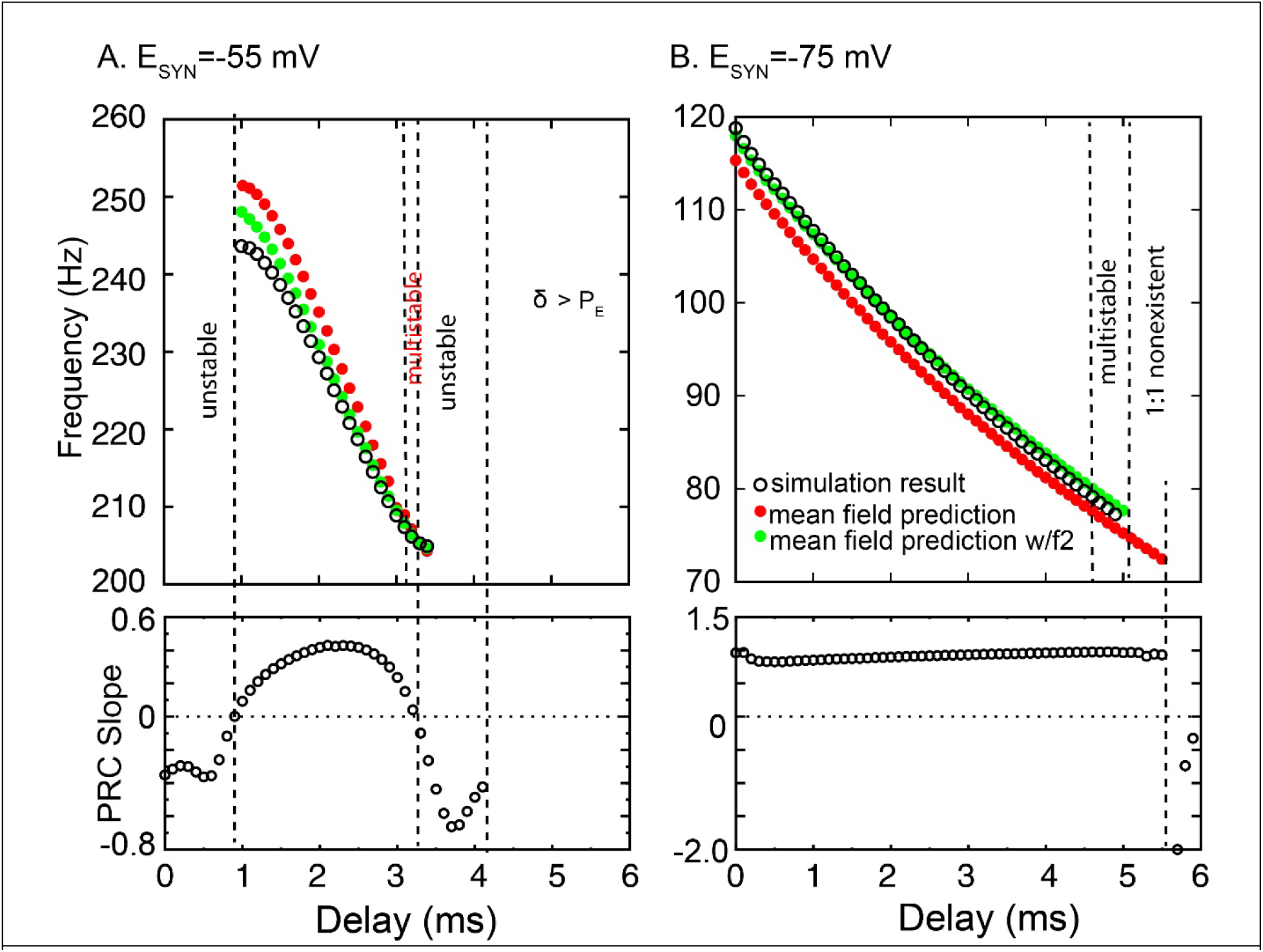
Comparison of Mean Field Predictions of 1:1 Synchrony with Observed Values from Simulations. Top predicted network frequency at each delay. Bottom; First order PRC slope at a given delay with the predicted network frequency. A. Shunting Inhibition. B. Hyperpolarizing Inhibition.

A major assumption of our analysis thus far is that a synchronous self-locking mode exists at *P_F_ =P_E_(1+f_1_(δ/P_E_))*, whether it is stable or not. However, we have previously shown that second order resetting can break this assumption [7]. Therefore, we plotted the second order mean field phase resetting for the individual oscillator in Fig. 5A and B1 for shunting and hyperpolarizing inhibition respectively. Note that in general the second order resetting has to the opposite sign as first order resetting (compare to Fig. 2C). The discontinuity in the first order PRC remarked upon in the text corresponding to Fig. 2 is also manifested in the second order resetting curve in Fig. 5B1, and for the same reason. The bulk of the area under the curve of the conductance waveform falls in cycle P2, hence the large values of second order resetting at late phases in Fig. 5B1. In addition to the existence criterion described in Fig. 3B, there is another existence criterion for global synchrony. The map of the intervals in Fig. 3A can be reinterpreted such that the stimulus interval *ts* contains the second order resetting. For 1:1 global synchrony, *ts* =δ. Previously we defined the phase at which an input is received as φ= δ/P_E_. To account for second order resetting, we redefine that phase as φ+ f_2_(φ) = δ/P_E._ This criterion again lends itself to a graphical method illustrated in Fig. 5B2. The delay values for 5.2-5.6 ms fall in the discontinuity, hence self-locking in a 1:1 mode cannot exist. Instead, we observed a 2:2 locking both in the 100 neuron network (not shown) and in a single neuron self-locked with N=36 inputs (Fig. 5C). The analysis assuming a biexponential train as Fig. 1 does not apply in this case because the inputs (bottom trace) are period two, with two distinct intervals characterizing this mode. The first bump in the conductance waveform is the delayed self-input due to the first spike. It occurs too late to prevent the next spike (see vertical dashed line). The delayed input from the second spike comes slightly more than 5.2 ms after the first one. They summate and produce a very long interval, then the sequence repeats. This 2:2 self-locking explains the strange hybrid modes that were observed to be multistable with global synchrony: some cycles produce 1:1 locking and other 2:2 lockings (Fig. 5D).

**Figure 5.**
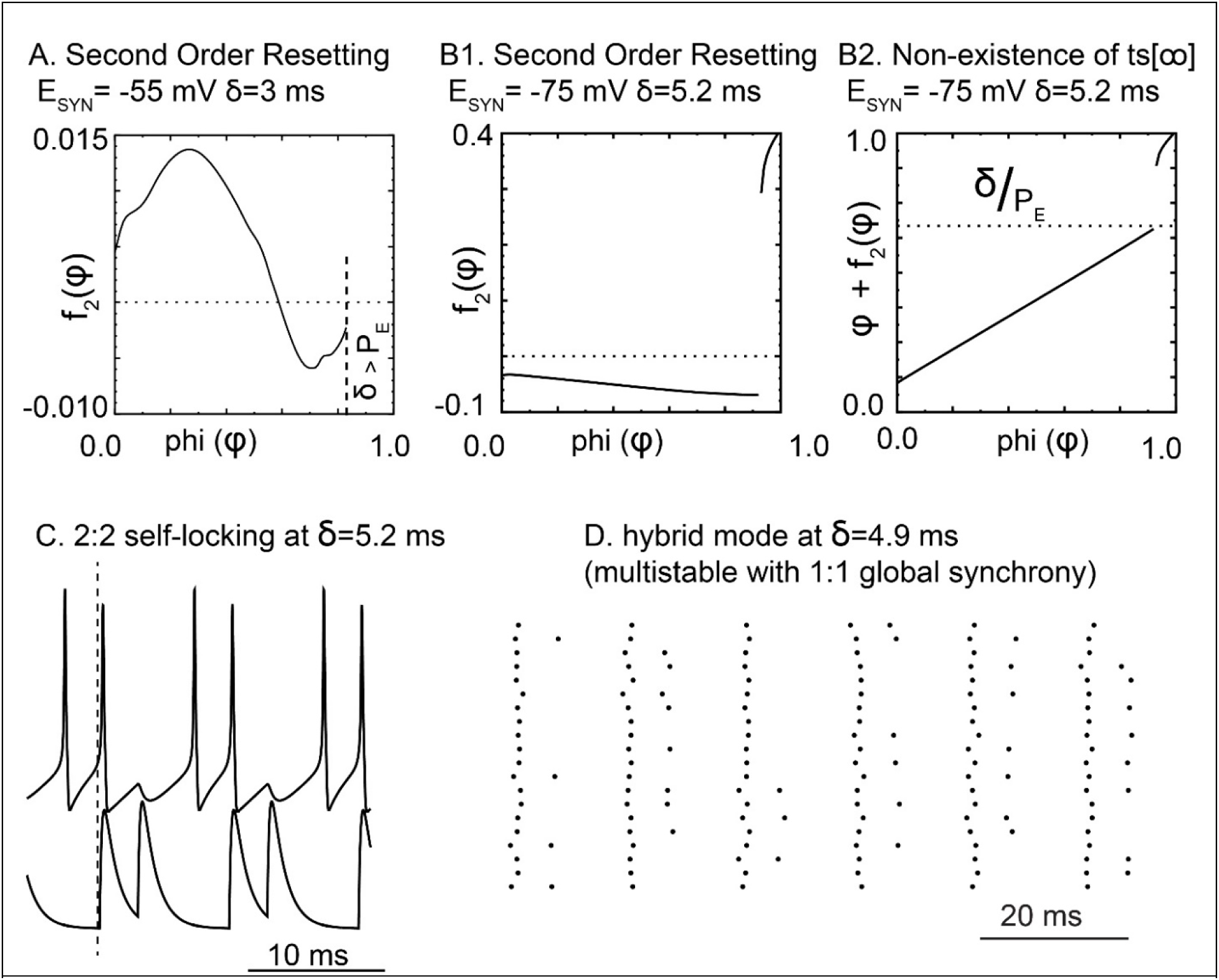
Second Order Resetting. A. Second order PRC for shunting inhibition. B1. Second order PRC for hyperpolarizing inhibition. B2. Existence criterion for stimulus interval based on second order resetting. C. Simulated self-connected cluster 2:2 locking. Top: voltage trace, bottom: conductance of self-connection. D. Downsampled raster plot of 20 of the 100 neurons in the network during a hybrid nonsynchronous mode.

After adding an existence criterion based on second order resetting, we updated the original criterion from Fig. 3 to include second order resetting: *P_F_=P_E_(1+f_1_(δ/P_E_) + f_2_(δ/P_E_)).* The plots in Fig. 3B actually reflect this criterion. We revisited our predictions of the network frequency in Fig. 4A and B to include second order resetting (green filled circles). The green filled circles nearly eliminate the systematic errors in prediction of the observed frequency. Moreover, the second order existence criterion mostly removes the incorrect predictions of 1:1 synchrony for delays from 5.1 to 5.6 ms in Fig. 4B.

## 4 Discussion

### Extending the applicability of pulsatile coupling methods

One of the major limitations of pulsatile coupling methods [4] has been that realistic synapses are not instantaneous. The literature on phase response curves has focused on resetting that occurs in the oscillatory cycle that contains the perturbation, called first order resetting. Methods to incorporate second order resetting that occurs in the cycle immediately following the cycle that contains the perturbation have been developed [46,47] and are utilized herein. However, the discrete maps upon which these methods are based cannot accommodate third and higher resetting. The mean field method presented in this paper allows for all results based on first (and where necessary second) order resetting to be generalized in a way that no longer requires the resetting due to one input be complete before the next input arrives, simply by substituting the embedded period for the free-running intrinsic period in the stability criteria as we did in Eq. 2.4. In this study, we based our stability predictions on the firing pattern in which a single oscillator is perturbed from global synchrony, and the predictions were in very good agreement with observations from simulated networks. Other perturbations from global synchrony [52,54,55] could also be used to predict the stability of global synchrony, likely producing similar predictions for stability of global synchrony as shown in [7].

A different approach to the summation of multiple inputs considered the case in which an oscillator does not return to its original limit cycle by the time the next input is received [56] by explicitly generating the PRC for multiple inputs. To some extent, this work is a generalization of that study.

### Second Order Resetting

We added second order resetting to the existence criterion in this study, but not to the criterion from stability of synchrony. If we adopt the terminology that *m_i,j_* is the slope of the *ith* order PRC for the *jth* oscillator, then per [6] the eigenvalues that determine stability are the cube roots of the following polynomial:

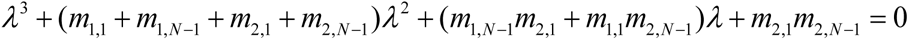

However, all terms are negligible in our case compared to the slope of the first order resetting of the single oscillator by the remainder of the network, *m_1,1_* in this notation, except in cases for hyperpolarizing inhibition in which stability did not exist anyway. Thus we did not need to consider second order resetting in our stability analysis; however, this may not be true generally.

### Caveat

Another limitation has been that neurons may possess slow processes that adapt in response to multiple inputs. Functional phase response curves [57], in which an input is applied repetitively and time locked to spike emission in neural oscillators, has been proposed as a method for understanding synchronization of neurons that exhibit adaptation. The mean field method does not account for adaptation, so it does not replace the functional phase response method approach to adaptation.

### Stochastic population oscillations versus coupled oscillators

A popular theory for fast oscillations in networks of inhibitory neurons [58–60] is based on waxing and waning waves of inhibition that are a network phenomenon that do not depend upon the phase resetting properties of individual neurons. This mechanism for stochastic population oscillations was developed to conform to the prevailing opinion that when the local field potential exhibits coherent fast oscillations, for example sharp-wave ripples, “the spike trains of constituent neurons are typically irregular and sparse. The dichotomy between rhythmic local field and stochastic spike trains presents a challenge to the theory of brain rhythms in the framework of coupled oscillators [61]”. It has long been noted that interneurons do not necessarily fire sparsely [33] during fast oscillations. As stated in the Introduction, numerous studies show that parvalbumin positive fast spiking interneurons (PV-INs) are phase-locked to ripple cycles and often fire at ripple frequency both *in vivo* [19–26] and *in vitro* [27–31]. Moreover, a very recent study [62] found that during ripples in area CA1 *in vivo*, PV FS interneurons form assemblies that spike synchronously within narrow time windows, providing strong support for a coupled oscillator mechanism, although the relative contributions to synchrony of synaptic inhibition between interneurons versus electrical connectivity or synaptic excitation from PV cells are not clear from that study.

### Conclusion

Although the theoretical method presented herein only strictly applies to homogeneous networks, we and others have demonstrated that the synchronization tendencies exhibited by the homogeneous networks studied here can persist even when substantial perturbations are introduced in the form of intrinsic heterogeneity and sparse connectivity [42,60,63]. Moreover, PV-INs in general are connected by electrical synapses [41,64–66], neglected here but which serve to mitigate heterogeneity [42]. The potential biological relevance of this work is that cognitive defects resulting from poor synchronization of fast oscillations might be ameliorated by manipulations that promote synchrony. The intrinsic phase response properties of inhibitory interneurons are likely essential to the understanding of fast oscillations and provide putative therapeutic targets for cognitive deficits that depend upon intact oscillatory substrates.

## 5 Computational Methods

Parameters for python code for the homogeneous network of parvalbumin positive inhibitory interneurons were taken from Fig. 4 and 5 of [42]: g_Na_ 16805 nS, g_Kv_1__ 59 nS, g_Kv_3__ 631.710 nS, E_L_ -72 mV and C_M_ 0.0768 nF. g_L_ was 14.7 nS resulting in an input resistance of 68 MΩ, and g_ChR_ was 7 nS, with others as in Table 1.Connections between fast spiking PV neurons in the medial entorhinal cortex have been well-established [41]although one study inexplicably failed to find them [62]. Each neuron received exactly 36 chemical synapses with a strength of 1.65 nS. The synaptic delay was varied.

**Table 1.** Parameters for gating variables. The gating parameters for the homogeneous network.

| | $m$ | $h$ | $n$ | $a$ |
| --- | --- | --- | --- | --- |
| $\theta$ | -53.0 | -55.71 | 5.9 | 51.36 |
| $\sigma_1$ (mV) | 4 | -20 | 12 | 12 |
| $\sigma_2$ (mV) | -13 | 3.5 | -8.5 | -80 |
| $k_1$ (ms<sup>-1</sup>) | 0.25 | 0.012 | 1 | 1 |
| $k_2$ (ms<sup>-1</sup>) | 0.1 | 0.2 | 0.001 | 0.02 |

